# Conidial fusion in the asexual fungus *Verticillium dahliae*

**DOI:** 10.1101/2020.12.16.423040

**Authors:** Vasileios Vangalis, Michael Knop, Milton A. Typas, Ioannis A. Papaioannou

**Affiliations:** Department of Genetics & Biotechnology, Faculty of Biology, National and Kapodistrian University of Athens, Greece; Center for Molecular Biology of Heidelberg University (ZMBH), Heidelberg, Germany; German Cancer Research Center (DKFZ), DKFZ-ZMBH Alliance, Heidelberg, Germany

**Keywords:** anastomosis, Conidial Anastomosis Tube (CAT), dikaryon, heterokaryon, nuclear degradation, parasexual cycle

## Abstract

Cell-to-cell fusion is a fundamental biological process across the tree of life. In filamentous fungi, somatic fusion (or anastomosis) is required for the normal development of their syncytial hyphal networks, and it can initiate non-sexual genetic exchange processes, such as horizontal genetic transfer and the parasexual cycle. Although these could be important drivers of the evolution of asexual fungi, this remains a largely unexplored possibility due to the lack of suitable resources for their study in these puzzling organisms. In this study, we report that the spores of the important asexual plant-pathogenic fungus *Verticillium dahliae* often engage in cell fusion via Conidial Anastomosis Tubes (CATs). We optimized appropriate procedures for their highly reproducible quantification and live-cell imaging, which were used to characterize their physiology and cell biology, and to start elucidating their underlying genetic machinery. Formation of CATs was shown to depend on growth conditions and require functional Fus3 and Slt2 MAP kinases, as well as the NADPH oxidase NoxA, whereas the GPCR Ste2 and the mating-type protein MAT1-2-1 were dispensable. We show that nuclei and other organelles can migrate through CATs, which often leads to the formation of transient dikaryons. Their nuclei have possible windows of opportunity for genetic interaction before degradation of one by a presumably homeostatic mechanism. We establish here CAT-mediated fusion in *V. dahliae* as an experimentally convenient system for the cytological analysis of fungal non-sexual genetic interactions. We expect that it will facilitate the dissection of sexual alternatives in asexual fungi.

## Introduction

Cell-to-cell fusion is a biological process of fundamental importance across the tree of life. Gamete fusion during fertilization is an integral component of sexual life cycles, which generate genetic diversity in eukaryotes (Brukman et al. 2019). Non-gametic cell fusion is also vital in multicellular animals and plants for the development of complex tissues and organs (Maruyama et al. 2016; Brukman et al. 2019). It is also a process of particular significance to filamentous fungi. These typically exist as interconnected networks of branched tubular cells called hyphae (Mela et al. 2020). During growth, hyphae engage in cell communication, which can lead to hyphal fusion, also known as anastomosis, within or between colonies. This involves chemotropic attraction and physical contact of cells, followed by localized dissolution of cell walls, plasma membrane merger and, eventually, establishment of cytoplasmic continuity through fusion bridges that connect different hyphae (Hickey et al. 2002; Fischer and Glass 2019). Frequent hyphal fusion leads to the formation of the characteristic hyphal syncytium that radiates by polarized tip elongation to colonize its substrate (Riquelme et al. 2018). Hyphal fusion reinforces these syncytia structurally and ensures rapid transport of nutrients throughout the colony (Hickey et al. 2002; Simonin et al. 2012; Roper et al. 2015).

Anastomosis also occurs at the earlier stages of fungal development, between conidial germlings (i.e. germinated asexual spores), via so-called Conidial Anastomosis Tubes (CATs) (Roca et al. 2005b). Germling fusion increases the chances of successful substrate colonization by permitting the efficient use of limited or heterogeneously distributed resources (Roca et al. 2005b; Mela et al. 2020). Similarly to hyphal fusion, CAT-mediated germling fusion appears widespread in fungi (Roca et al. 2005b). The process of germling fusion is considered mechanistically equivalent to and under the same genetic control as hyphal fusion in mature colonies (Fu et al. 2011; Fleißner and Herzog 2016).

When anastomosis takes place between colonies or germlings of different genotypes (i.e. non-self-fusion), heterokaryons may arise, where genetically distinct nuclei share the common cytoplasm of the fused cells (Mela et al. 2020). This can expose these syncytial organisms to the rapid spread of selfish genetic elements (e.g. mycoviruses and transposable elements), and therefore heterokaryon formation and viability are strictly regulated and limited by a number of allorecognition checkpoint systems (Gonçalves and Glass 2020). Despite those, however, non-self somatic fusion and heterokaryon formation have been implicated in several cases in the horizontal transfer of genetic material, such as lineage-specific chromosomes (Soanes and Richards 2014; Vlaardingerbroek et al. 2016). In addition, they can initiate parasexual processes of global genetic exchange and recombination under selective pressure in the lab. Parasexuality involves nuclear fusion in heterokaryons and restoration of the original ploidy level by random chromosome loss. During this process, random reassortment of chromosomes and frequent mitotic crossing over contribute to the generation of novel genetic combinations in the progeny, thereby promoting genetic diversity (Webster 1974; Sherwood and Bennett 2009; Strom and Bushley 2016). Although the contribution of parasexual cycles in the evolution of filamentous fungi has not yet been clarified, parasexuality has been proposed as an ancestral process in the mitosis-to-meiosis evolutionary trajectory (Goodenough and Heitman 2014), with potential significance in extant fungi such as yeasts (Anderson et al. 2019; Zhao et al. 2020).

Our current understanding of the mechanisms underlying hyphal and germling communication and fusion are largely based on the extensively studied sexual ascomycete *Neurospora crassa* (Roca et al. 2005a; Fischer and Glass 2019; Gonçalves and Glass 2020). In addition, the cell biology of CATs has been described, to some extent, in the species *Colletotrichum lindemuthianum* (Roca et al. 2003), *Botrytis cinerea* (Roca et al. 2012), *Fusarium oxysporum* (Kurian et al. 2018) and *Zymoseptoria tritici* (Francisco et al. 2020). With the only exception of *F. oxysporum*, these fungi have well-established sexual reproduction cycles and use meiotic recombination to reshuffle their genomes and create genetic variation. We envision that in organisms that lack sexual reproduction, such as the asexual fungi (Dyer and Kück 2017), somatic fusion could be an important driver of evolution by granting them access to genetic interactions that could substitute for sexual reproduction to generate diversity. However, this remains a largely unexplored possibility, since our understanding of cell communication and fusion is almost exclusively based on sexual species, mostly *N. crassa*, in which the parasexual cycle is of limited or no relevance. Therefore, the dissection of these processes in suitable asexual representatives could provide valuable insights into fungal genetics and evolution.

A particularly interesting candidate is the widespread and economically important plant pathogen *Verticillium dahliae*. A sexual stage of this fungus has never been identified and multiple lines of evidence (i.e. extreme mating-type bias, frequent karyotypic polymorphisms and clonal expansion) consistently suggest lack of sexual activity (Papaioannou et al. 2013b; de Jonge et al. 2013; Gurung et al. 2014; Short et al. 2014). However, population analyses have detected considerable levels of recombination, with the potential to create new lineages in this species (Atallah et al. 2010; Milgroom et al. 2014). We hypothesize that asexual genetic processes, such as horizontal genetic transfer or parasexuality, could be the source of recombination and explain this conundrum. A full parasexual cycle has long been demonstrated in *Verticillium* species and used in genetic analyses (Hastie 1964; Puhalla and Mayfield 1974; Typas and Heale 1976; Typas 1983). Remarkably, *Verticillium* species were found to have special parasexual features, including unusually high instability of heterozygous diploids and extremely high frequencies of mitotic recombination (Hastie 1967, 1968; Typas and Heale 1978). These considerations suggest that *V. dahliae* could be a particularly useful organism for the elucidation of parasexual mechanisms and their possible contribution to fungal evolution.

In this study, we aimed at the characterization of somatic cell fusion in *V. dahliae*, the first and essential step that could provide the species with access to non-sexual genetic exchange. For this, we focused on the identification of CAT-mediated cell fusion, we analyzed its dependence on growth parameters and genetic factors, and we optimized experimental procedures for its reproducible quantification and convenient live-cell imaging. We expect that the establishment of CAT-mediated cell fusion in *V. dahliae* as an experimentally feasible and consistent system for the cytological analysis of genetic phenomena will significantly contribute to the future research of the genetics and evolution of this and other important asexual fungi.

## Results

### Conidial Anastomosis Tube (CAT)-mediated cell fusion in *V. dahliae*

Our preliminary investigations of somatic fusion in *V. dahliae* aimed at the characterization of hyphal fusions in mature colonies of the species (Fig. 1a). However, the high hyphal density and complexity of the mycelial networks rendered the microscopic quantification of hyphal fusion a formidable task. We therefore decided to focus on the study of fusion at the stage of conidial germlings, which is known to be equivalent to fusion of mature hyphae. Our first attempts to identify conidial fusion in *V. dahliae* were hampered by their strong inhibition by nutrients in all standard media (Fig. 2a). However, when we incubated conidia of our reference strain (123V) in water for 3 days, numerous anastomoses were systematically observed between conidia and/or conidial germlings and hyphae (Fig. 1b). Under these conditions, conidia started to germinate after ∼ 20 h, while the first anastomosis bridges were detected ∼ 12 h later. Both processes appeared dependent on the conidial adhesion to the glass or polystyrene substrate of the culture plates, which generally took place before germination and/or fusion. The observed bridges were significantly thinner than germ tubes (average widths: 1.58 ± 0.18 μm and 2.35 ± 0.21 μm, respectively; n = 150 each; independent samples *t*-test, *p*-value: 1.4 × 10^−4^), and they exhibited an average length of 6.67 ± 4.12 μm (n = 150). Their maximum detected length was 25.88 μm. In a sample of 1,000 such bridges, they were most frequently formed between germlings (33.8%) or between ungerminated conidia (30.4%), and less frequently between a germling and an ungerminated conidium (19.0%) or between a germling and a hypha (11.8%). More rarely, bridges were identified between an ungerminated conidium and a hypha (3.8%), while secondary connections between already established bridges were also observed (1.2%) (Fig. 1b). Interconnected networks of conidia and germlings with multiple anastomoses were often observed (Fig. 1b; movie S1).

**Fig. 1.**
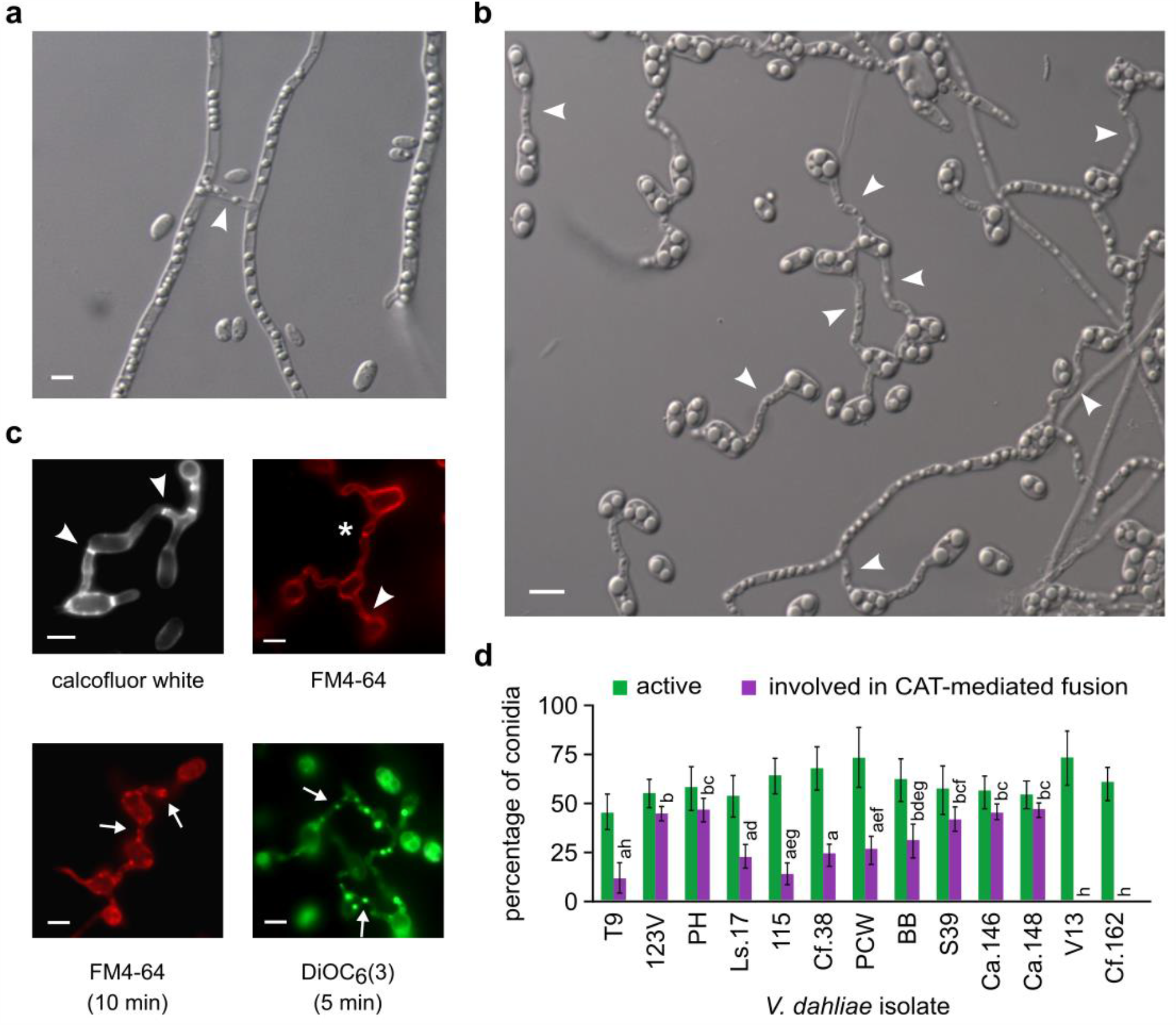
Somatic cell-to-cell fusion in *V. dahliae*. **a** Anastomosis (arrowhead) between mature hyphae of isolate 123V grown on water agar. **b** Anastomoses between conidia, germlings and hyphae of isolate 123V, following incubation in water (pH 7.6) for 3 days. Representative examples of different types of CATs are marked with arrowheads. **c** Fluorescence microscopy images of representative CATs of isolate 123V stained with calcofluor white, FM4-64 or DiOC_6_(3). Imaging was performed immediately after the addition of the dye (top row) or following incubation of the duration specified in brackets (bottom row). Arrowheads: septae within CATs; asterisk: incompletely formed CAT at the stage before plasma membrane merger (evidenced by the typical constriction of the CAT that surrounds the FM4-64 signal); arrows: examples of organelles/vesicles migrating through CATs. Bars (in **a**-**c**) = 5 μm. **d** Frequencies of active conidia (i.e. conidia losing their original cell symmetry by germinating and/or being involved in fusion) and active conidia participating in CAT-mediated fusion events, of *V. dahliae* isolates of all VCGs. Each isolate was tested in triplicate, and 300 conidia were analyzed per replicate. Bars = SD. Statistical significance of differences was tested by one-way ANOVA, followed by Tukey’s post-hoc test (bars with the same letter are not significantly different; *p* ≤ 0.05).

**Fig. 2.**
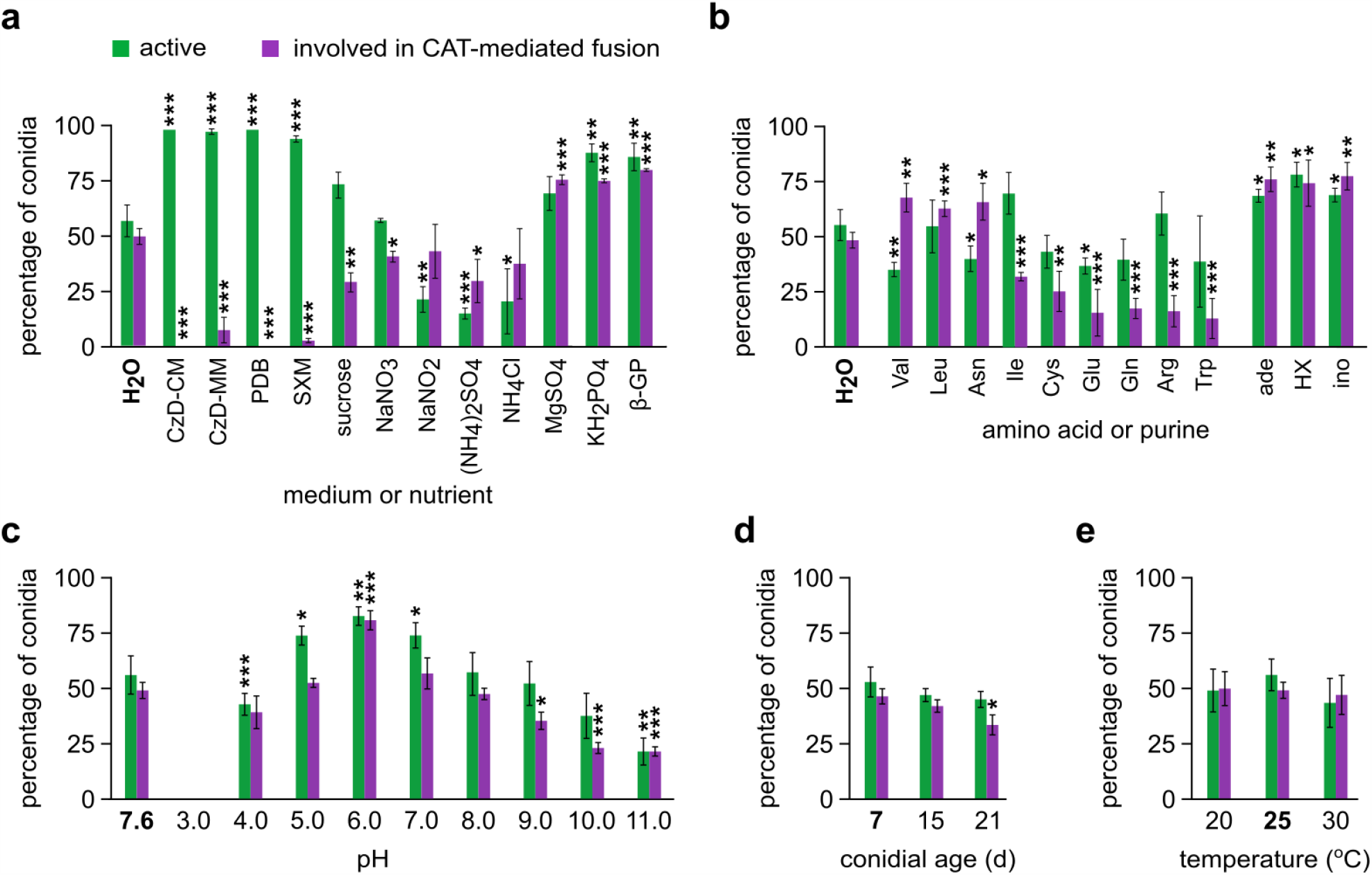
Influence of growth medium or nutrient (**a**), amino acid or purine (**b**), pH (**c**), conidial age (**d**) and temperature (**e**) on the frequencies of active conidia and active conidia involved in CAT-mediated fusion (*V. dahliae* isolate 123V). Each medium/condition was tested in triplicate, and 300 conidia were analyzed per replicate. Bars = SD. CzD: Czapek-Dox medium; SXM: simulated xylem sap medium; β-GP: β-glycerophosphate; HX: hypoxanthine; ino: inosine. Statistical significance of differences from the control (in bold characters in each plot) was tested by Student’s *t*-test (* *p* ≤ 0.05, ** *p* ≤ 0.01, *** *p* ≤ 0.001). The results from the remaining amino acids are presented in Fig. S1.

At least one septum was identified in 68.0% of the anastomosis bridges (n = 150), as we determined by using calcofluor white for fluorescent staining of chitin in cell walls (Fig. 1c). Furthermore, we used FM4-64 and DiOC_6_(3) for staining of organellar membranes (i.e. endocytic compartments and endoplasmic reticulum, mitochondria and vesicles, respectively). Fluorescence microscopy often suggested translocation of organelles through such anastomoses (Fig. 1c). Overall, the observed anastomosis bridges in *V. dahliae* are similar to the Conidial Anastomosis Tubes (CATs) previously characterized in other fungi (Roca et al. 2005b), and they will be referred to as such hereinafter.

### Frequency of CAT formation varies among isolates and depends on growth conditions and conidial age

To test whether CAT formation levels in *V. dahliae* are isolate-dependent, we compared the behavior of 13 wild-type isolates from various hosts, geographic origins and all major Vegetative Compatibility Groups (VCGs) of the species (Table S1). Isolates of all VCGs readily formed CATs at frequencies that ranged between 12.1-50.5% of active conidia (i.e. conidia losing their original cell symmetry by germinating and/or being involved in fusion), depending on the isolate (Fig. 1d). The only two isolates that did not form any CATs were V13 and Cf.162, which had previously been characterized as heterokaryon self-incompatible (Papaioannou et al. 2014; Papaioannou and Typas 2015).

Apart from the strong inhibition of CAT formation by the nutrient-rich media PDB and Czapek-Dox supplemented with rich nutrient sources (CzD-CM), the frequency of CATs was still diminished when conidia were incubated in the defined Czapek-Dox minimal medium (MM) (Fig. 2a). By testing separately the effect of the ingredients of this medium and other related substances on CAT formation, we observed a significant inhibitory effect of the carbon source (sucrose) and modest inhibitory effects of the nitrogen sources NaNO_3_, NaNO_2_, (NH_4_)_2_SO_4_ and NH_4_Cl. In contrast to sucrose and NaNO_3_, which did not affect germination negatively, the other four substances significantly compromised germination, which is probably responsible for the observed reductions of CAT levels in these cases. On the other hand, MgSO_4_, KH_2_PO_4_ and β-glycerophosphate significantly promoted CAT levels. These findings indicate that the strong inhibition imposed by CzD-MM is due to combined effects and/or the overall levels of several nutrients rather than to the dominant effect of any particular ingredient. Furthermore, CAT formation was detectable, albeit at a very low frequency, in a xylem sap-simulating medium (SXM), which opens up the possibility of its occurrence *in planta* during the infection cycle.

We also tested the effects of all standard amino acids and purines (or their derivatives) on CAT formation (Fig. 2b and Fig. S1). We found that valine, leucine and asparagine clearly promoted CAT formation, in contrast to isoleucine, cysteine, glutamic acid, glutamine, arginine and tryptophan, which reduced CAT levels with only small or no effects on conidial germination (Fig. 2b). No significant effects were exerted on CAT formation by the remaining amino acids (Fig. S1). In the presence of the purine adenine or its derivatives hypoxanthine and inosine, both germination and CAT formation were significantly increased (Fig. 2b).

The optimal pH for CAT formation was 6.0, which lies in the middle of the optimal range 5.0-7.0 for germination (Fig. 1c). Frequency of CATs was symmetrically reduced in both acidic and alkaline directions in the pH range 4.0-8.0, and CAT formation was still possible at lower levels up to pH 11.0. Conidial age showed a minor influence on CAT competence, with conidia from older colonies being less proficient in fusion (Fig. 1d). Temperature of incubation had no effect on CAT formation in the range from 20 to 30 °C (Fig. 1e). Finally, in the course of our experiments we observed that CAT levels were dependent on conidial concentration. We determined as optimal ranges and routinely used in our study 5.0 × 10^6^ - 1.0 × 10^7^ conidia/mL in wells of standard 24-well cell culture plates, and 5.0 × 10^5^ - 1.0 × 10^6^ conidia/mL in 96-well plates.

### Types of CAT-mediated fusion determined by time-lapse live-cell imaging

To gain a better understanding of the CAT formation process in *V. dahliae*, we optimized an experimentally convenient method for live-cell imaging of anastomosis. We used this procedure in time-lapse imaging experiments of *V. dahliae* interacting germlings that engaged in CAT fusion. In these experiments, we studied both self-fusion between isogenic germlings (isolate Ls.17) and non-self-fusion between germlings of isolates Ls.17 and Cf.38, which belong to the same VCG (Table S1). For the identification of the isolates, we labeled their nuclei with different fluorescent proteins. In particular, we constructed strains expressing histone H1-mCherry or H1-sGFP in the background of isolate Ls.17, and a Cf.38 derivative strain expressing H1-sGFP. Time-lapse imaging data of a total of 204 CAT-mediated fusion events were recorded and analyzed (i.e. 96 self-fusion and 108 non-self-fusion events).

The whole process of CAT formation, from homing to fusion, lasts 20 to 180 min, during which nuclei of the interacting cells never divide (Fig. 3). Our investigation revealed two morphologically distinct types of CAT formation. The first fusion type (type I) involves the protrusion of a CAT from one of the interacting germlings, its attraction and growth towards its partner and, finally, its fusion with a “peg” that is induced in the latter shortly before fusion (Fig. 3a; movie S2). In the second case (type II), both interacting cells form tubes that, upon a phase of growth orientation towards each other (homing), eventually come into physical contact and fuse (Fig. 3b; movie S3). Type II was more frequent in both self- and non-self-fusions (56.3% and 60.2%, respectively), with no significant difference between the two pairings (*χ*^*2*^-test; *p* > 0.05). Conidial anastomosis tubes often emerged from the tips of germ tubes (Fig. 3b; movie S4).

**Fig. 3.**
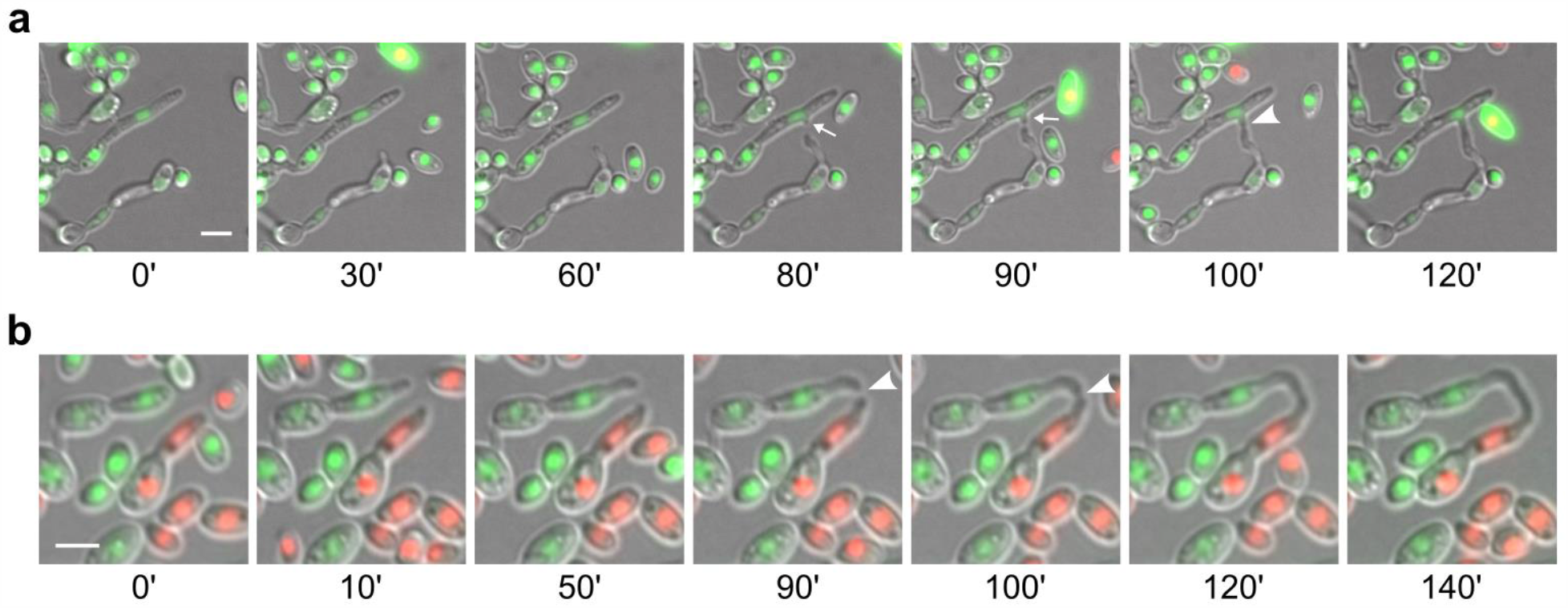
Types of CAT formation in *V. dahliae*, as revealed by live-cell time-lapse imaging. **a** Self-fusion in strain Ls.17 expressing H1-sGFP for nuclear labeling. A CAT is initiated from one of the interacting cells and is finally fused to an induced “peg” (arrows) of the other. **b** Non-self-fusion between strains Ls.17 (expressing H1-mCherry) and Cf.38 (H1-sGFP). Symmetrical tip-to-tip attraction and CAT fusion between the germ tubes of the two partners. Arrowheads: contact point. Bars = 5 μm.

### CAT-mediated dikaryon formation and nuclear migration

The hypothesis that vegetative fusion could initiate the parasexual cycle in *Verticillium* species relies on the assumption that fusion is completed upon physical contact of hyphae or germlings, and that cytoplasmic continuity is established between the fused cells. We examined this in *V. dahliae* directly using live-cell imaging. To this end, we constructed a strain expressing cytoplasmic sGFP in the background of isolate Ls.17 and incubated its conidia with non-fluorescent conidia of either the same isolate (self-pairing) or of isolate Cf.38 (non-self-pairing) (Table S1). We then performed time-lapse imaging of 35 self- and 12 non-self-fusions. Our experiments clearly demonstrated that CAT formation in both pairings always led to the establishment of cytoplasmic continuity between the interacting cells. This manifested itself microscopically as a sudden burst of rapid cytoplasmic transfer between the two partner cells ∼ 20-30 min after their physical contact. This was followed in all cases by a phase of slower motion and gradual intermixing of cytoplasmic material (Fig. 4a; movie S5). No differences were detected in the dynamics of this process between self- and non-self-pairings. Notably, we often observed temporary restriction of cytoplasmic flow within the CATs or the recipient young hyphae, at the septae that compartmentalized them (Fig. 4a). This is consistent with the well-established roles of the septal pore in the control of cytoplasmic transfer between the otherwise continuous cells. Our data provides direct support for the possibility of dikaryon formation as the result of cell fusion in *V. dahliae*, an essential prerequisite for the occurrence of the parasexual cycle.

**Fig. 4.**
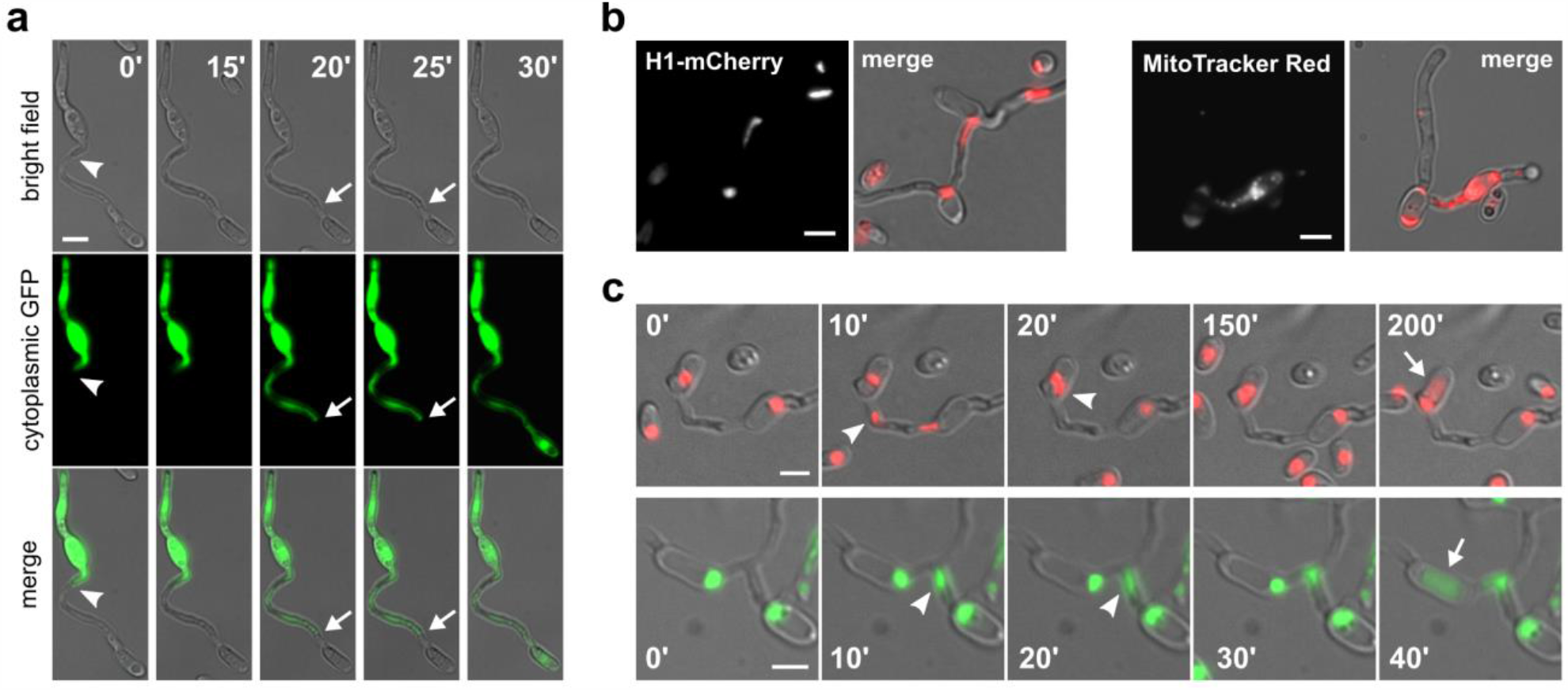
Establishment of cytoplasmic continuity and migration of genetic material through CATs in *V. dahliae*. **a** Time-lapse imaging of CAT formation between the wild-type isolate Ls.17 and a Ls.17 derivative strain expressing cytoplasmic sGFP. At time point 0’ the interacting tubes have established physical contact (arrowhead) but not yet membrane merger, as evidenced by the characteristic constriction around the contact point and the lack of cytoplasmic transfer. A septum in the CAT is indicated by arrows. **b** Migration of a nucleus (in strain Ls.17 H1-mCherry) and mitochondria (strain Ls.17, staining with MitoTracker Red) through CATs. **c** Time-lapse imaging of migrating nuclei (arrowheads) through CATs, in strain Ls.17 engineered to express H1-mCherry or H1-sGFP for nuclear labeling. Nuclear translocation was followed by degradation (arrows) in the recipient cell. Bars = 5 μm.

These findings served as our motivation to further investigate the possibility of transfer of genetic material through CATs, which would be needed for the direct fusion of haploid nuclei to diploid formation. The first evidence for transfer of genetic material through CATs was obtained by fluorescence microscopy of the isolate Ls.17 after staining its mitochondria with MitoTracker Red, and of the engineered Ls.17 H1-mCherry strain with labeled nuclei. We recorded many instances of both mitochondria and elongated nuclei entering CATs or apparently migrating through CATs (Fig. 4b). We further performed time-lapse imaging of the Ls.17 H1-mCherry and Ls.17 H1-sGFP strains (a total of 85 anastomoses were analyzed) to prove this definitively. Nuclear translocation through CATs was observed in 34 of them (40.0%), in both strains (Fig. 4c). This occurred 0.5-2.0 h after dikaryon formation, and it always followed division of one of the two nuclei of the dikaryon. Migration of one of the daughter nuclei always started soon after the completion of mitosis (in less than 5 min, which was the time resolution of most of our time-lapse data), and it was usually complete within 10-20 min. Translocation of nuclei through septae in CATs was never observed, indicating that those CATs that develop a septum experience a translocation-competent phase of limited duration, which is completed with the septum formation. These findings are in agreement with our previous observations of nuclear migration through septal pores being extremely rare in *V. dahliae* (Vangalis et al. 2020).

Following nuclear transfer, two nuclei co-existed for some time in the recipient compartment, usually in close proximity to each other (Fig. 4c). However, direct nuclear fusion was never observed in our study. In some cases, degradation of one of the two nuclei was observed after a phase of dikaryotic co-existence, always leaving the other nucleus apparently unaffected. Degradation was highly unsynchronized, starting 10 min to 3 hours after the completion of nuclear translocation. Although we did not observe this phenomenon in all cases, it is currently unclear whether it affects only a fraction of the fused cells or whether it happens always but with a timing that could sometimes exceed the maximum duration of our imaging movies.

### Conserved signaling and cytoskeletal components are involved in CAT-mediated fusion

We next asked whether the molecular mechanisms that govern cell-to-cell fusion in the asexual fungus *V. dahliae* are similar to the ones that have been described in the sexual model species *N. crassa* and other fungi. To investigate this, we focused on conserved components with important functions in signal transduction. We also asked whether cytoskeletal components are important for CAT-mediated fusion in *V. dahliae*. To address these questions, we first tested the effects of a number of inhibitors of signaling pathways and the cytoskeleton on CAT formation, to gain some first insights into their hypothesized contributions to the process (Fig. 5a). Remarkably, inhibitors of polymerization of both microtubules (oryzalin) and the actin cytoskeleton (cytochalasin B, ML-7) drastically reduced CAT formation, in contrast to germination that appeared to depend only on microtubules. Regarding signaling, an inhibitor of phospholipase D activity (1-butanol) also affected severely conidial competence for CAT formation, while a similar but less pronounced effect was also observed on germination. For Mitogen-Activated Protein Kinase (MAPK) signaling, we used UO126 (inhibits the mammalian MAPKKs MEK1/2, which phosphorylate p42/44, and could thus interfere with fungal MAPK cascades of Fus3 and Slt2), and SB203580 (inhibits the p38 MAPK, homolog of the Hog1 kinase of fungi). Of these two inhibitors, the former specifically diminished CAT frequency (no effect on germination), in contrast to the latter that had no effect on CAT formation. Finally, the inhibitor of the Reactive Oxygen Species (ROS)-generating NADPH oxidases (DPI) significantly prevented CATs without exhibiting any effect on germination. Therefore, functional microtubules and actin seem to be necessary for CAT-mediated fusion, as well as wild type-level activities of phospholipase D and NADPH oxidase, and functional MAP kinase cascades, except for the hyperosmotic glycerol (HOG) pathway.

**Fig. 5.**
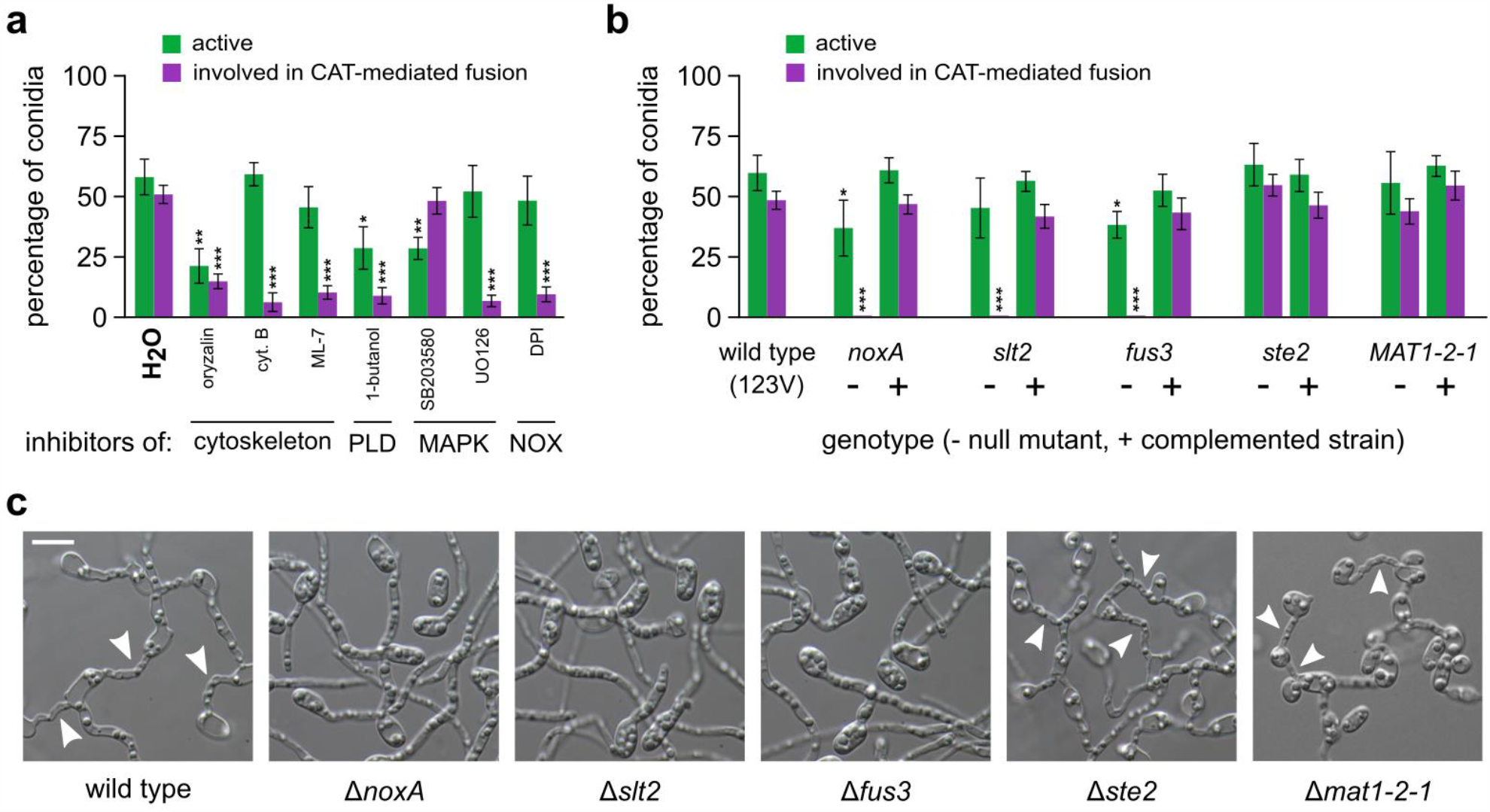
Somatic cell fusion in *V. dahliae* requires conserved signaling and cytoskeletal components. **a** Influence of inhibitors of cytoskeletal elements and the signaling components phospholipase D (PLD), MAP kinase cascades (MAPK), and ROS-generating NADPH complexes (NOX) on the frequencies of active conidia and active conidia involved in CAT-mediated fusion (isolate 123V). **b** Analysis of the fusion ability of *V. dahliae* 123V knock-out mutants for selected genes of NOX signaling (*noxA*), MAPK signaling (*slt2, fus3* and *ste2*) and the mating-type gene (*MAT1-2-1*). Frequencies of active conidia and active conidia involved in CAT-mediated fusion are shown for each null mutant (-), in comparison to their corresponding rescued strains by complementation (+), and the wild type. Each inhibitor and mutant was tested in triplicate, and 300 conidia were analyzed per replicate. Bars = SD. Statistical significance of differences from the control (H_2_O or the wild type, respectively) was assessed using Student’s *t*-tests (* *p* ≤ 0.05, ** *p* ≤ 0.01, *** *p* ≤ 0.001). **c** Fusion ability of the null mutants in comparison to their wild-type strain (*V. dahliae* 123V).

Based on this evidence, which suggests the involvement of particular MAPK and ROS signaling components in the control of CAT-mediated fusion in *V. dahliae*, we decided to further elucidate these functional links by constructing null mutants of *V. dahliae* 123V for selected genes that are involved in MAP kinase signaling (Δ*fus3*, Δ*ste2* and Δ*slt2*) and ROS generation (Δ*noxA*). *Fus3* and *ste2* in the model yeast species *S. cerevisiae* code for the MAP kinase that is involved in mating, and the G protein-coupled receptor (GPCR) of the alpha-factor pheromone that acts upstream of the Fus3 cascade, respectively. *Slt2*, on the other hand, codes in yeast for the MAP kinase of the cell wall integrity pathway. *NoxA* codes for the conserved in eukaryotes (though absent from yeast) superoxide-generating NADPH oxidase A, a major cellular source of ROS. Finally, we decided to also test the *V. dahliae* mating-type gene *MAT1-2-1*, of unknown function in this asexual organism. We first identified single homologs of these conserved genes in the *V. dahliae* genome using BLAST searches (VDAG_09461 for *fus3*, similarity to the *Sc*Fus3 protein: 78%, E-value = 9.0 × 10^−155^; VDAG_05622 for *ste2*, similarity to the *Fusarium oxysporum* Ste2 protein: 57%, E-value = 2.0 × 10^−67^; VDAG_02584 for *slt2*, similarity to the *Sc*Slt2 protein: 81%, E-value = 0; VDAG_06812 for *noxA*, similarity to the *F. oxysporum* NoxA protein: 92%, E-value = 0; and VDAG_02444 for *MAT1-2-1*, similarity to the *F. oxysporum* MAT1-2-1 protein: 55%, E-value = 6.0 × 10^−26^). We then knocked them out individually in *V. dahliae* 123V by gene disruption (*Vdfus3*; Fig. S2a) or deletion (all the others; Fig. S2b), using protoplast or *Agrobacterium tumefaciens*-mediated transformation, respectively. Transformants were screened for homologous integration by PCR (Fig. S2c) and verified by Southern blot analyses (Fig. S2d). Complemented strains were generated for all genes by re-introducing into the knock-out strains the corresponding wild-type genes (open reading frames), flanked by ∼ 2.0 kb-long genomic sequences that presumably contained the native promoters and terminators of the genes.

When the knock-out strains were tested for their competence in CAT-mediated fusion, mutants Δ*fus3*, Δ*slt2* and Δ*noxA* were reproducibly shown to be totally unable to form CATs, even under optimal conditions (Fig. 5b). Their germination patterns were only moderately affected in comparison to the wild type, and complementation with the corresponding wild-type alleles always rescued their phenotypes. On the other hand, mutants Δ*ste2* and Δ*mat1-2-1* retained the ability to germinate and form CATs at wild-type levels. These results clearly indicate that functionality of both Fus3 and Slt2 MAP kinase pathways is essential for somatic cell fusion in *V. dahliae*. However, the requirement for Fus3 is uncoupled from Ste2, which is the upstream activator of the Fus3 MAPK cascade in *S. cerevisiae*. In addition, ROS generation by NADPH oxidase A is also indispensable for fusion between *V. dahliae* conidia. The mating-type gene is not required for anastomosis.

## Discussion

Generation of genetic diversity is critical for the evolution of organisms and it represents a major function of sexual reproduction, a ubiquitous feature of eukaryotic life cycles (Goodenough and Heitman 2014). Despite the benefits of sex, though, approximately 20% of all fungi are thought to lack a sexual cycle. While many of those could be retaining some “cryptic” or conditional potential for sex (Dyer and Kück 2017), several lines of evidence suggest that non-sexual alternatives for genetic exchange and reshuffling might play a role in the enigmatic evolution of some of these organisms. Such alternatives could be the horizontal genetic transfer between isolates or species (Soanes and Richards 2014; Vlaardingerbroek et al. 2016), and parasexuality, possibly an ancestral but also a derived state in fungi (Pontecorvo 1956; Goodenough and Heitman 2014; Anderson et al. 2019; Zhao et al. 2020). The occurrence of such genetic processes in filamentous fungi largely depends on somatic cell fusion and subsequent dikaryon formation, which are very little understood in asexual fungi. Therefore, we undertook this study to optimize suitable experimental procedures for the analysis of CAT-mediated cell fusion in the asexual plant-pathogenic fungus *V. dahliae*. We used them to describe the physiology and cell biology of CATs in this species, and to start unraveling its genetics. We expect that these will prove essential resources for the successful elucidation of possible non-sexual genetic processes in this and other important fungi.

Given the inherent difficulties involved in the microscopic quantification of hyphal fusion in complex mycelia of mature colonies, we focused on CAT-mediated fusion of germlings. This process is mechanistically similar to hyphal fusion and has been shown to be under the same genetic control in *N. crassa* (Fu et al. 2011; Fleißner and Herzog 2016). The study and analysis of CATs proved to be much more convenient, requiring only straightforward experimental setups and, most importantly, permitting highly reproducible quantification and consistent live-cell imaging of fusion. It allowed uncomplicated fluorescence microscopy experiments for the characterization of CATs, as well as the time-lapse analysis of CAT formation and the nuclear fate in the resulting dikaryons. Quantification of CAT-mediated fusion also facilitated the analysis of the effects on cell fusion of inhibitory agents targeting cytoskeletal and signaling components, and the screening of mutants compromised in somatic fusion.

The anastomosis bridges between *V. dahliae* germlings showed many typical characteristics of fungal CATs, including emergence from germinated but also from ungerminated conidia, positive chemotropism (homing), significantly smaller width than germ tubes, determined growth potential (the longest CAT detected in our study was approximately 25 μm-long), development of septae in the majority, dependence on conidial density and pH, and requirement for both microtubule and actin cytoskeleton, in contrast to germination that requires only functional microtubules. All of these features have been reported in at least some of the other studied species (Roca et al. 2003, 2005a, 2012; Kurian et al. 2018; Francisco et al. 2020). This indicates that somatic cell fusion in fungi has conserved underlying core mechanisms and components among phylogenetically distant species with diverse lifestyles, regardless of their ability to undergo sexual reproduction. We investigated this notion further by testing the involvement of conserved signaling components of fungal somatic fusion in CAT-mediated fusion in *V. dahliae*. We found that both Fus3- and Slt2-homologous MAP kinases, as well as NADPH oxidase A are essential for fusion in *V. dahliae*, similarly to other tested fungi (Prados Rosales and Di Pietro 2008; Roca et al. 2012; Kayano et al. 2013; Dirschnabel et al. 2014; Segorbe et al. 2017; Fischer and Glass 2019; Nordzieke et al. 2019; Francisco et al. 2020). However, the requirement for *Vd*Fus3 is uncoupled from *Vd*Ste2, which is not needed for CAT formation, in agreement with studies of *N. crassa* (Kim and Borkovich 2004). Similarly, the mating-type protein *Vd*MAT1-2-1 is dispensable for conidial fusion. Inhibition of phospholipase D also compromised the ability of *V. dahliae* to form CATs, and this has also been observed in two distant species (Hassing et al. 2020). Overall, these findings underline that somatic fusion is mediated, indeed, by conserved signaling pathways in ascomycetes, which are distinct from the mechanisms underlying gametic fusion.

Anastomosis of conidia is severely inhibited in *V. dahliae* by the presence of nutrients in the growth medium. This is consistent with similar observations in the plant pathogens *C. lindemuthianum* (Ishikawa et al. 2010), *F. oxysporum* (Kurian et al. 2018) and *Z. tritici* (Francisco et al. 2020), but in contrast to the saprophyte *N. crassa*, which is competent at CAT-mediated fusion even in nutrient-rich media (Fischer-Harman et al. 2012). This difference could reflect the adaptation of plant pathogens to the challenging conditions that they face during infection, which presumably render the optimization of nutrient use critical for their success (Divon and Fluhr 2007). This hypothesis is further supported by our following findings: i. the inhibition of fusion is not due to any particular ingredient of our standard media, but to the overall levels of nutrients or the combined effect of multiple ingredients, and ii. CAT formation in *V. dahliae* is positively or negatively affected by several amino acids, which is also different from the situation in *N. crassa*, where CAT-mediated fusion is sensitive only to tryptophan (Fischer-Harman et al. 2012). Overall, our data suggests that the regulation of CAT formation differs between *V. dahliae* and *N. crassa* in that in the former it is more tightly linked to the environmental availability of nutrients. Notably, CAT formation in *V. dahliae* was also possible in a plant xylem sap-simulating medium, albeit at a very low frequency, which nevertheless suggests that it might be possible *in planta*. This might be of particular significance in the case of co-infection of a plant by different *V. dahliae* isolates, as it opens up the possibility of non-sexual genetic interactions between them within the plant tissues.

Conidial germling fusion via CATs was detected, at varying but significant frequencies of up to approximately 50% of active conidia, in *V. dahliae* isolates of all VCGs, but not in the heterokaryon self-incompatible isolates V13 and Cf.162. Such isolates fail to establish prototrophic heterokaryons when their auxotrophic mutants are paired with complementary auxotrophic mutants of the same or other isolates (Papaioannou et al. 2014; Papaioannou and Typas 2015). It is reasonable to assume that their general inability to form heterokaryons directly results from their observed defects in anastomosis. This link suggests that cell-to-cell fusion may indeed be essential for the initiation of genetic exchange processes between genetically dissimilar individuals. The first requirement for this would be the establishment of cytoplasmic continuity between the fused cells through the connecting CATs. This was clearly demonstrated in our experiments, both in self- and non-self-interactions of germlings, which showed that fusion leads to the formation of dikaryons, in which nuclei share the common cytoplasm of stably anastomosed cells. We further studied the behavior of nuclei in such “self” dikaryons to discover that nuclear migration through a CAT occurred very often, within 2 h after the completion of fusion. We have also made similar observations, though at different frequencies, in “non-self” dikaryons (our unpublished results). Following nuclear transfer, the two nuclei co-existed in the same compartment, usually in close physical proximity (no direct fusion of nuclei was observed, though), for up to several hours, until the end of our imaging movies. This was followed, in some cases, by selective degradation of one nucleus only, while the other remained apparently unaffected. As this phenomenon is quite unsynchronized, it is not clear whether it affects a fraction of dikaryons or if it is the general fate of these nuclei, which however might take in some cases longer than the duration of our time-lapse movies to begin.

Similar phenomena of nuclear degradation following transfer through anastomosis bridges have also been observed in other fungi (Ruiz-Roldán et al. 2010; Ishikawa et al. 2012), but their biological function is elusive. Given that degradation affects only one nucleus, while the other survives in an otherwise viable cell, and because it is also observed in self-pairings (i.e. isogenic nuclei), we conclude that it does not represent a typical heterokaryon incompatibility mechanism. These are triggered by genetic differences of the two nuclei and would typically result either in the prevention of fusion or in the destruction of the heterokaryotic cell (Gonçalves and Glass 2020). It is possible that in fungi such as *V. dahliae*, which almost exclusively have one nucleus per cellular compartment and nuclei cannot normally migrate through septal pores (Vangalis et al. 2020), the observed degradation of the additional nucleus in the dikaryon might be the result of a homeostatic mechanism, possibly involving autophagy (Corral-Ramos et al. 2015), that strictly regulates the number of nuclei per cell. In any case, however, one can speculate that even if nuclear degradation is a general phenomenon, the co-existence of nuclei for up to several hours before the onset of degradation, and the presence of chromatin of the degraded nucleus in the cell after nuclear disorganization provide possible windows of opportunity for genetic interactions of the two nuclei, as has been previously discussed (He et al. 1998; Richards et al. 2011; Vlaardingerbroek et al. 2016). These could include horizontal transfer of chromatin, or even fast nuclear division and fusion while the nuclei are in physical contact with each other. These exciting questions particularly welcome future research for the elucidation of genomic and organismic evolution in the absence of sex, and the understanding of the genetics and population dynamics of the enigmatic asexual fungi.

## Materials and methods

### Fungal isolates and VCG classification, growth media and culture conditions

Representative wild-type *V. dahliae* isolates of all major Vegetative Compatibility Groups (VCGs), isolated from different hosts and geographic origins, were included in this study and are listed in Table S1. The vegetative compatibility behavior of all isolates has been extensively characterized before (Papaioannou et al. 2014; Papaioannou and Typas 2015), with the exception of isolate 123V, which was tested in this study and classified into VCG 2A. For this, we used our previously described procedures to generate nitrate non-utilizing (*nit*) mutants, classify them into *nit1* and *nitM* groups, and use them in complementation tests with *nit* mutants of tester strains of all VCGs (Papaioannou and Typas 2015). Each pairing was performed in triplicate. Standard complete media (Potato Dextrose Agar - PDA, Czapek-Dox complete medium - CM) and growth conditions (24 °C, in the dark) were used for growth of fungal strains; preparation and maintenance of monoconidial strains have been previously described (Papaioannou et al. 2013b).

### Conidial anastomosis (CAT) assays and optimization

Anastomosis of conidia/conidial germlings by formation of CATs was routinely assayed in wells of either polystyrene 24-well cell culture plates (662160 from Greiner Bio-One, Kremsmünster, Austria) or glass-bottom 96-well plates (MGB096-1-2-LG-L from Matrical Bioscience, Spokane, WA, USA), which were directly used for live-cell time-lapse fluorescence microscopy. For each CAT formation assay, conidial suspensions were freshly prepared in water from 7-day-old PDA cultures of the appropriate isolates, and their concentrations were determined using a Neubauer improved cell counting chamber. Conidia were then transferred to wells of cell culture plates, at final concentrations in the range 5.0 × 10^6^ - 1.0 × 10^7^ conidia/mL for 24-well cell culture plates, and 5.0 × 10^5^ - 1.0 × 10^6^ conidia/mL for 96-well plates. In the case of non-self-pairings, 1:1 mixtures of conidial suspensions of the appropriate isolates were used, at the same final concentrations. Plates were statically incubated at 24 °C (in the dark) for 72 h, before imaging. Each assay was performed in at least three independent repetitions.

The same batch of autoclaved water was used in all CAT-mediated fusion assays, with pH 7.6 and the following chemical composition (mg/L): total hardness (CaCO_3_) 219, HCO_3_^-^ 244, Cl^-^ 4.29, SO_4_^2-^ 9.16, NO_3_^-^N 1.93, F^+^ 0.09, Ca^2+^ 83, Mg^2+^ 3.06, Na^+^ 2.85 and K^+^ 1.02 (heavy metals undetectable). The Potato Dextrose Broth (PDB) tested during the optimization of CAT assays was purchased from Scharlab (02-483; Barcelona, Spain). Our modified Czapek-Dox minimal medium (MM) had the following composition (g/L): sucrose 30.0, KH_2_PO_4_ 0.35, KCl 0.5, NaNO_3_ 2.0, KCl 0.5, FeSO_4_ · 7H_2_O 0.033, MgSO_4_ · 7H_2_O 0.62, β-glycerophosphate disodium salt · 5H_2_O 0.765 (pH 6.8). We used the same Czapek-Dox medium, with the addition of yeast extract, malt extract, casein hydrolysate and bacteriological peptone, each at 2.0 g/L (pH 6.8) as a nutrient-rich Czapek-Dox complete medium (CM). We also tested a previously described simulated xylem fluid medium (SXM), which resembles the nutritional conditions *in planta* (Neumann and Dobinson 2003).

The ingredients of our modified Czapek-Dox MM were also tested individually in CAT formation assays, each at the same concentration as the one used in the medium. Other tested nutrients and mineral sources were used at the following concentrations (μg/mL): NaNO_2_ 50; (NH_4_)_2_SO_4_ 200; NH_4_Cl 200. All amino acids and purines were used at the final concentration of 2 μg/mL. Inhibitors of cytoskeletal elements and conserved signaling components were used as follows: oryzalin 40 μM, cytochalasin B 50 μg/mL, ML-7 20 μM, 1-butanol 0.2%, SB203580 50 μM, UO126 20 μM, DPI 50 μM. All chemicals were purchased from Merck (Darmstadt, Germany) or Sigma-Aldrich (St. Louis, MO, USA).

### Microscopic determination of CAT-mediated fusion frequency

After the end of the incubation period, CAT formation assay samples were observed undisturbed under the microscope. For this, we used a Zeiss (Oberkochen, Germany) Axioplan microscope equipped with a differential interference contrast optical system and a Zeiss Axiocam MRc5 digital camera. At least three independent assays were performed for each tested isolate or condition and 300 conidia were analyzed per replicate. Discrimination between germ tubes and CATs at pre-fusion stages is not always possible, and conidial germination under starvation conditions is extremely unsynchronized. To overcome these difficulties and obtain reproducible and reliable comparisons between isolates and conditions, we consistently calculated for each sample: i. the percentage of “active” conidia as the fraction of conidia that had lost their cell symmetry by the end of the incubation period (this predominantly happens because of germination, but also emergence of CATs cannot be excluded); and ii. the percentage of conidia “involved in CAT-mediated fusion” as that fraction of active conidia that participated in at least one fusion event. These criteria permit meaningful comparisons between conditions as they take into account differences in conidial viability and germination kinetics, and their use is consistent with the previously published literature.

### Disruption or deletion of *V. dahliae* genes

Standard and previously described protocols were used for PCR amplification (a full list of PCR primers is provided in Table S2; all of them were obtained from Eurofins Genomics, Ebersberg, Germany or Sigma-Aldrich, St. Louis, MO, USA), bacterial transformation (*E. coli* strain DH5α), plasmid extraction, restriction digestion, Sanger sequencing (Eurofins Genomics) and maintenance (Papaioannou et al. 2013b, a). All plasmids constructed and used for fungal transformation are listed in Table S3. The hypervirulent *Agrobacterium tumefaciens* strain AGL-1 was transformed with appropriate binary vectors according to previously reported procedures (Vangalis et al. 2020). We followed our optimized and published protocols for protoplast and *A. tumefaciens*-mediated transformation of *V. dahliae* (Vangalis et al. 2020).

The experimental strategies that were followed for disruption (*Vdfus3*) or deletion (*Vdslt2, Vdste2, VdMAT1-2-1, VdnoxA*) of *V. dahliae* genes are schematically described in Fig. S2a-b. Briefly, for disruption of *Vdfus3*, a 0.5 kb-long internal fragment of *Vdfus3* was sub-cloned and ligated to vector pUCATPH (Table S3) to generate the disruption vector pUCfus3 (Fig. S2a). This was used to transform protoplasts of *V. dahliae* 123V. Deletion of the four other genes was achieved by transforming *V. dahliae* 123V (*A. tumefaciens*-mediated transformation) with the corresponding deletion vectors in the backbone of plasmid pOSCAR (Table S3, Fig. S2b). Upstream and downstream homologous arms (∼ 2.0 kb-long) of each gene were amplified from genomic DNA of *V. dahliae* and ligated to the flanking regions of the *hph* or *neo*^R^ resistance cassette (amplified from plasmids pUCATPH and pSD1, respectively; Table S3) to generate the deletion constructs between the left and right borders (LB and RB, respectively) of *A. tumefaciens* binary vector pOSCAR (Fig. S2b). The high-fidelity Herculase II Fusion DNA Polymerase (Agilent, Santa Clara, CA, United States) was used for PCR amplification. All transformants were screened by PCR using gene-specific primers (Table S2, Fig. S2c) and were verified by Southern hybridization analyses (DIG DNA Labeling and Detection Kit, Sigma-Aldrich, St. Louis, MO, USA) (Fig. S2d).

For complementation of the null mutants, the wild-type genes were re-introduced into the corresponding mutants. For this, their coding sequences and ∼ 2.0 kb-long flanking regions (presumably containing the native promoters and transcription terminators of the genes) were amplified from genomic DNA of *V. dahliae* 123V and co-transformed into protoplasts of the corresponding mutants with plasmids pUCATPH or pSD1 (containing the *hph* and *neo*^R^ cassettes that confer resistance to hygromycin B and geneticin, respectively).

### Nuclear labeling (histone H1) with mCherry or sGFP and cytoplasmic labeling with sGFP

For nuclear labeling with the fluorescent protein mCherry, we first used BLAST searches to identify the *V. dahliae* homolog of the histone H1 gene (VDAG_09854). We then used the KAPA HiFi DNA Polymerase (Roche, Basel, Switzerland) to amplify: i. a 1,940 bp-long PCR product from genomic DNA of *V. dahliae* isolate Ls.17 containing the open reading frame of *VdH1* (excluding the stop codon) and a 1,000 bp-long upstream fragment that presumably contains the native promoter of the gene; ii. a 3,203 bp-long amplicon from plasmid pAN8.1-mCherry (Ruiz-Roldán et al. 2010) containing the *mCherry* coding sequence and the phleomycin resistance (*ble*^R^) cassette; and iii. a 1,698 bp-long amplicon from *V. dahliae* Ls.17 genomic DNA presumably containing the *VdH1* transcription terminator sequence. The three amplicons were used for synthesis of a construct for the C-terminal tagging of VdH1 with mCherry using a fusion PCR strategy (Szewczyk et al. 2006). First, a PCR reaction was performed using the KAPA HiFi DNA Polymerase, 10 ng of each of the two amplified genomic fragments, and 50 ng of the glycine-alanine-*mCherry*-*ble*^R^ cassette, in the absence of PCR primers. The second PCR reaction was performed using 2.5 μL of the first PCR product as template, the primer pair VdH1FusF/VdH1FusR (Table S2) and the KAPA HiFi DNA Polymerase. The final fusion PCR product was used to transform protoplasts of *V. dahliae* Ls.17 for the C-terminal tagging of VdH1 with mCherry. For tagging of histone H1 with sGFP, we transformed protoplasts of *V. dahliae* isolates Ls.17 and Cf.38 with plasmid pMF357 (Table S3), which contains the *hph* cassette that confers resistance to hygromycin B, as well as a fusion construct of the *sGFP* gene to the *N. crassa* histone H1 gene. The *V. dahliae* Ls.17 strain with robust cytoplasmic expression of sGFP has been described before (Vangalis *et al*., 2021).

### Fluorescence microscopy and live-cell imaging

For staining of CATs with fluorescent dyes, conidia were first used in CAT formation assays as described above. At the end of the incubation period, each desired stain was added directly to the wells of the cell culture microplates, at the following final concentrations: calcofluor white M2R 10 μg/mL, DIOC_6_(3) 50 μM, FM4-64 25 μM and Mitotracker Red 10 μM (all from Sigma-Aldrich, St. Louis, MO, USA). Imaging was performed immediately or after incubation at room temperature for 5 min (calcofluor white M2R, DiOC_6_(3), and MitoTracker Red) or 10 min (FM4-64). A Zeiss (Oberkochen, Germany) Axioplan epifluorescence microscope was used, equipped with filter sets G 365 nm (excitation) and LP 420 nm (emission), BP 450-490 nm (excitation) and BP 515-595 nm (emission), and BP 510-560 nm (excitation) and LP590 nm (emission), as well as a Zeiss Axiocam MRc5 digital camera.

For live-cell time-lapse microscopy we used a Nikon (Tokyo, Japan) Ti-E epifluorescence microscope equipped with an autofocus system (Perfect Focus System, Nikon), a LED light engine (Spectra X from Lumencor, Beaverton, OR, USA), filter sets 469/35 and 525/50 or 542/27 and 600/52 (excitation and emission, respectively; all from Semrock, Rochester, NY, USA, except for 525/50, which was from Chroma Technology, Bellows Falls, VT, USA), and a sCMOS camera (Flash4.0 from Hamamatsu, Honshu, Japan). Images were acquired every 5 or 10 min for up to 24 h (exposure time: 50 ms for both the green and the red channel), and finally processed using ImageJ (Schindelin et al. 2012).

## Acknowledgements

We are grateful to M. P. Pantou for the Δ*fus3* mutant, which was constructed during her previous, unpublished work in the group of MAT (National and Kapodistrian University of Athens, Greece). Plasmid pAN8.1-mCherry was a generous gift from A. Di Pietro (University of Córdoba, Spain). This research is co-financed by Greece and the European Union (European Social Fund-ESF) through the Operational Programme «Human Resources Development, Education and Lifelong Learning» in the context of the project “Strengthening Human Resources Research Potential via Doctorate Research” (MIS-5000432), implemented by the State Scholarships Foundation (IKY).

## Author contributions

IAP and MAT conceived the study. IAP and VV planned and designed the experiments. VV and IAP performed the experiments, collected and analyzed the data. IAP wrote the manuscript, with input from VV and MAT. All authors commented on previous versions of the manuscript, read and approved the final version. MAT and MK provided access to funding and resources.

## Supplementary Figures

**Fig. S1.**
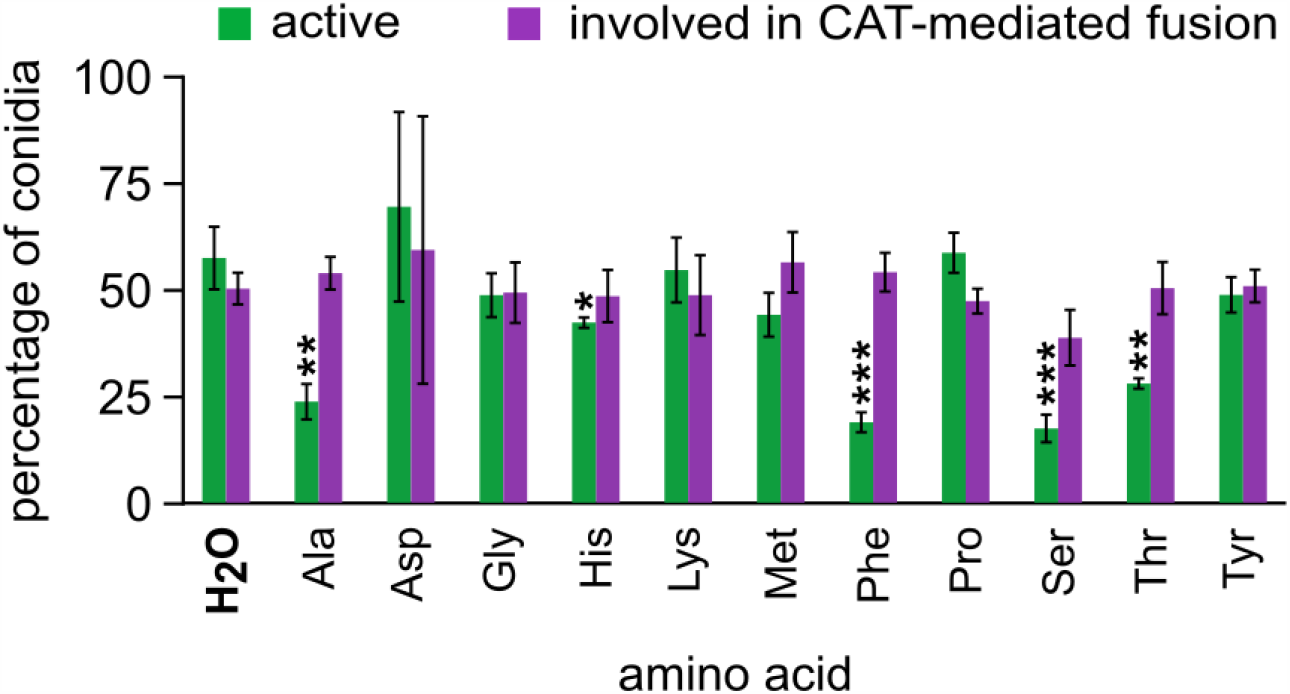
Influence of various amino acids on the frequencies of active conidia and active conidia involved in CAT-mediated fusion (*V. dahliae* isolate 123V). Each amino acid was tested in triplicate, and 300 conidia were analyzed per replicate. Bars = SD. Statistical significance of differences from the control was determined by Student’s *t*-test (* *p* ≤ 0.05, ** *p* ≤ 0.01, *** *p* ≤ 0.001). The results from the remaining amino acids are presented in Fig. 2a.

**Fig. S2.**
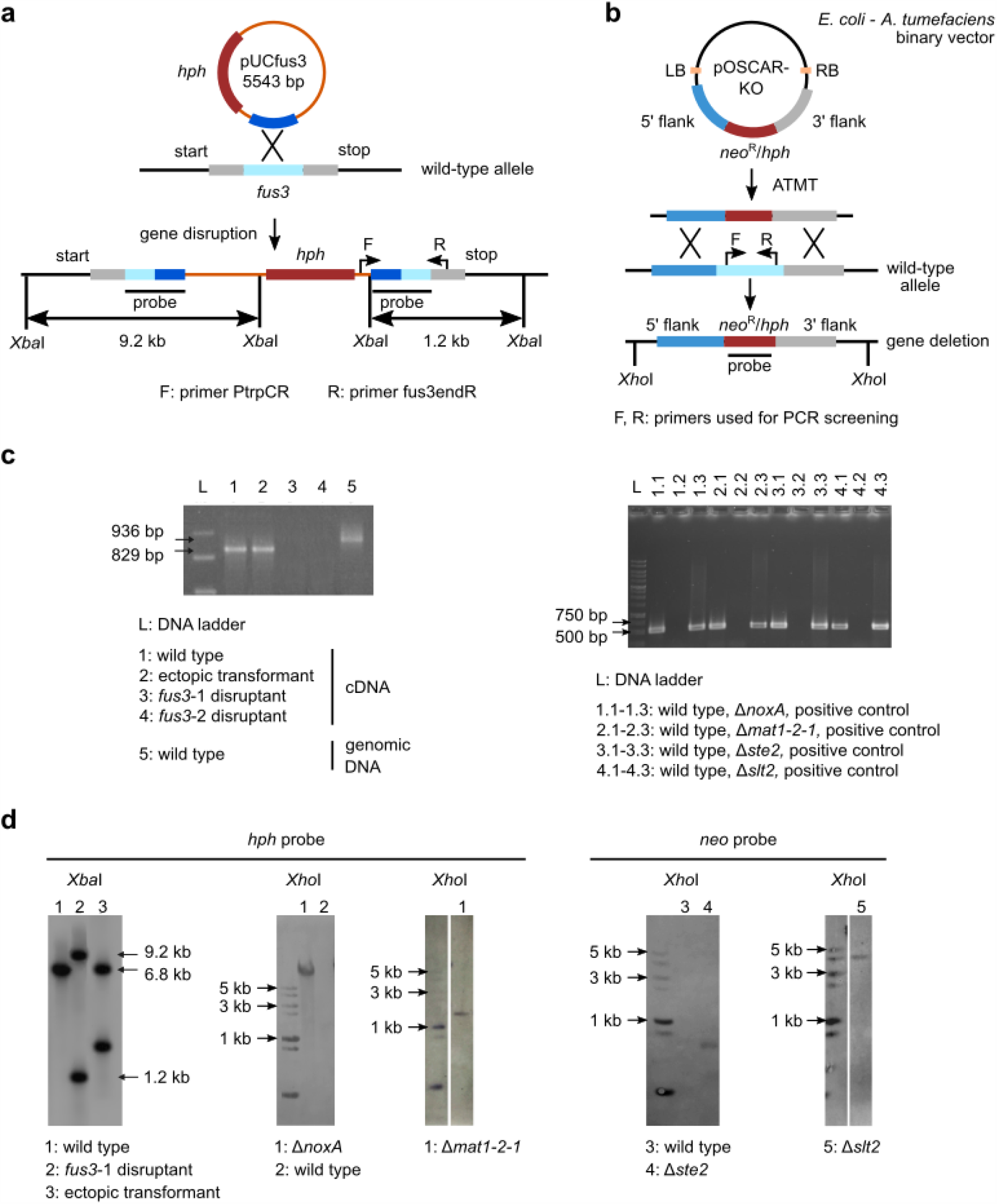
Construction and validation of null mutants used in this study. **a** Schematic representation of the gene disruption strategy that was followed for knocking out the *V. dahliae* homolog of *fus3*, using protoplast transformation of *V. dahliae* 123V. **b** A double homologous recombination-based strategy using *Agrobacterium tumefaciens*-mediated transformation was used for the deletion of the *V. dahliae* 123V homologs of genes *slt2, ste2, noxA* and *MAT1-2-1*. **c** PCR screening using gene-specific primers for disruption (*fus3*) or deletion (*slt2, ste2, noxA, MAT1-2-1*) mutants. **d** Validation of mutants by Southern hybridization experiments.

**Table S1.** *Verticillium dahliae* strains used in this study

| wild-type isolates |  |  |  |
| --- | --- | --- | --- |
| strain | VCG | host | origin |
| T9 | 1A | cotton | USA, CA |
| 123V | 2A<sup>1</sup> | tomato | Greece |
| PH | 2A | pistachio | USA, CA |
| Ls.17 | 2B | lettuce | USA, CA |
| 115 | 2B | cotton | Syria |
| Cf.38 | 2B<sup>2</sup> | chili pepper | USA, CA |
| PCW | 3 | pepper | USA, CA |
| BB | 4A | potato | USA, ID |
| S39 | 4B | potato | USA, OH |
| Ca.146 | 6 | bell pepper | USA, CA |
| Ca.148 | 6 | bell pepper | USA, CA |
| V13 | HSI<sup>3</sup> | cotton | Spain |
| Cf.162 | HSI<sup>2,3</sup> | chili pepper | USA, CA |

<sup>1</sup> The VCG of isolate 123V was determined in this study.
<sup>2</sup> Isolates Cf.38 and Cf.162 were originally misclassified in VCG 6. Their behavior was re-assessed and the VCGs provided here were determined by Papaioannou *et al.* (2014).
<sup>3</sup> HSI: Heterokaryon Self-Incompatible.

| constructed strains (this study) |  |  |  |
| --- | --- | --- | --- |
| strain | background | genotype | constructed with plasmid(s) |
| Ls.17-gfp | Ls.17 | *sgfp*, *neo*<sup>R</sup> | pIGPAPA |
| Ls-H1-mCherry | Ls.17 | *nit1*, *VdH1-mCherry*, *ble*<sup>R</sup> | fusion PCR product |
| Ls-H1-gfp | Ls.17 | *nitM*, *NcH1-sgfp*, *hph* | pMF357 |
| Cf.38-H1-gfp | Cf.38 | *nitM*, *NcH1-sgfp*, *hph* | pMF357 |
| 123-Δ*fus3* | 123V | *fus3::hph* | pUC*fus3* |
| 123-Δ*fus3*-c | 123-Δ*fus3* | *fus3::hph*, *fus3*, *neo*<sup>R</sup> | pUC*fus3*, *fus3*, pSD1 |
| 123-Δ*noxA* | 123V | *ΔnoxA::hph* | pOSCAR-*noxA* |
| 123-Δ*noxA*-c | 123-Δ*noxA* | *ΔnoxA::hph*, *noxA*, *neo*<sup>R</sup> | pOSCAR-*noxA*, *noxA*, pSD1 |
| 123-Δ*mat1-2-1* | 123V | *Δmat1-2-1::hph* | pOSCAR-*mat* |
| 123-Δ*mat1-2-1*-c | 123-Δ*mat1-2-1* | *Δmat1-2-1::hph*, *mat1-2-1*, *neo*<sup>R</sup> | pOSCAR-*mat*, *mat1-2-1*, pSD1 |
| 123-Δ*slt2* | 123V | *Δslt2::neo*<sup>R</sup> | pOSCAR-*slt2* |
| 123-Δ*slt2*-c | 123-Δ*slt2* | *Δslt2::neo*, *slt2*, *hph* | pOSCAR-*slt2*, *slt2*, pUCATPH |
| 123-Δ*ste2* | 123V | *Δste2::neo*<sup>R</sup> | pOSCAR-*ste2* |
| 123-Δ*ste2*-c | 123-Δ*ste2* | *Δste2::neo*, *ste2*, *neo*<sup>R</sup> | pOSCAR-*ste2*, *ste2*, pUCATPH |

**Table S2.**
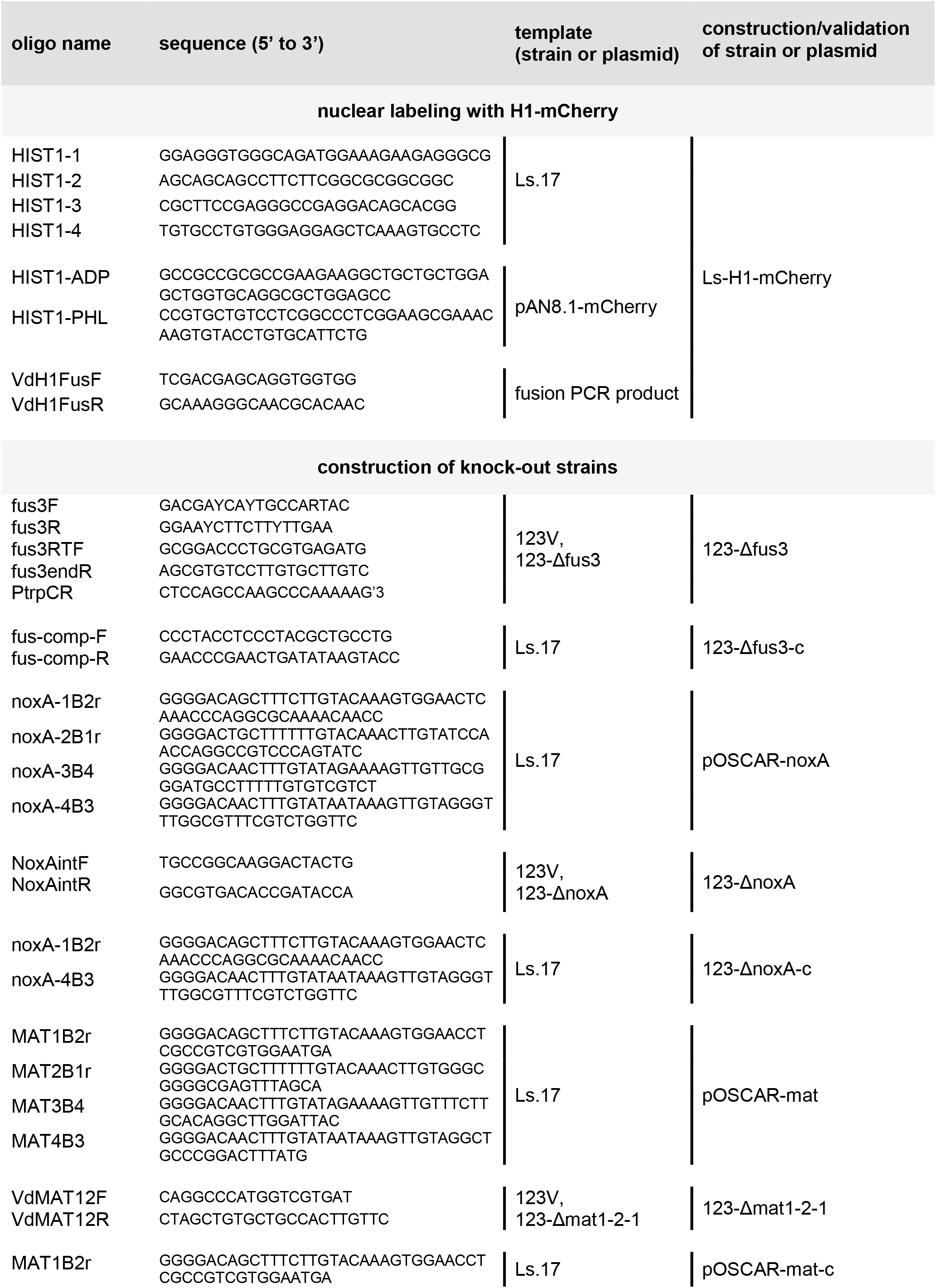
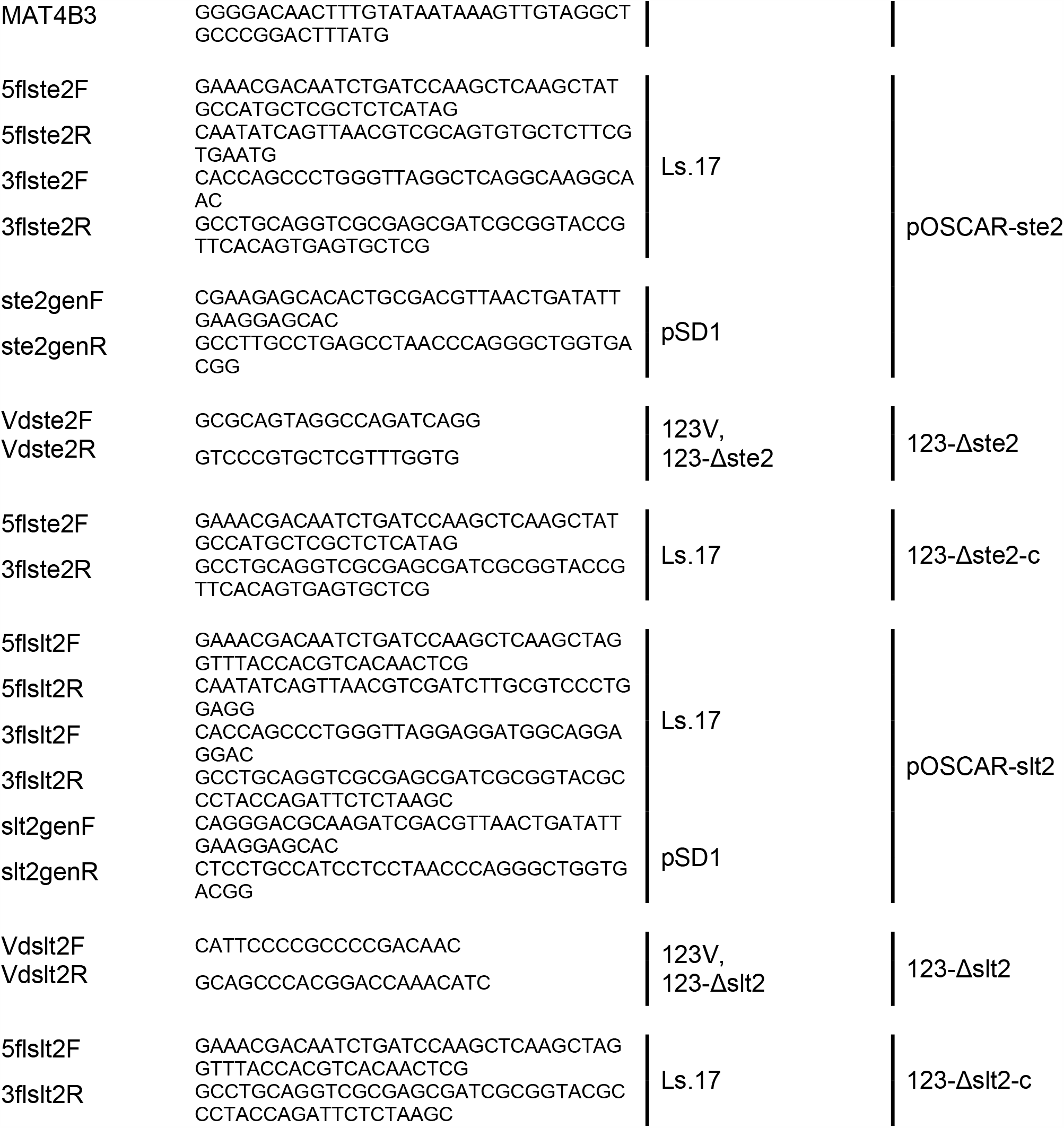
List of DNA oligonucleotides used in this study

**Table S3.** Plasmids constructed and used in this study

| plasmid | backbone | description | source |
| --- | --- | --- | --- |
| pIGPAPA | - | $P_{icr}$-*sgfp*-T<sub>nosI</sub>, *hph* | Horwitz *et al.*, 1999 |
| pUCATPH | - | *hph* | Lu *et al.*, 1994 |
| pSD1 | pBluescript II | $P_{gpdA}$, $P_{trpC}$, *neo*<sup>R</sup> | Nguyen *et al.*, 2008 |
| pOSCAR | pPZP-RCS2 | *A. tumefaciens* binary vector | Paz *et al.*, 2011 |
| pA-Hyg-OSCAR | - | *hph* | Paz *et al.*, 2011 |
| pAN8.1-mCherry | pAN8.1 | GA-*mCherryFP*, *ble*<sup>R</sup> | Ruiz-Roldan *et al.*, 2010 |
| pMF357 | - | $P_{ccg1}$-*NcH1-sgfp*, *hph* | Ishikawa *et al.*, 2012 |
| pOSCAR-mat | pOSCAR | 5'H<sub>Vdmat</sub>-*hph*-3'H<sub>Vdmat</sub><br>(H: 2.0 kb-long homology arms) | This study |
| pOSCAR-noxA | pOSCAR | 5'H<sub>VdnoxA</sub>-*hph*-3'H<sub>VdnoxA</sub><br>(H: 2.0 kb-long homology arms) | This study |
| pOSCAR-slt2 | pOSCAR | 5'H<sub>VdsIt2</sub>-*neo*<sup>R</sup>-3'H<sub>VdsIt2</sub><br>(H: 2.0 kb-long homology arms) | This study |
| pOSCAR-ste2 | pOSCAR | 5'H<sub>Vdste2</sub>-*neo*<sup>R</sup>-3'H<sub>Vdste2</sub><br>(H: 2.0 kb-long homology arms) | This study |
| pUCfus3 | pUCATPH | *fus3*, *hph* | This study |

